# The human acetylcholinesterase c-terminal T30 peptide activates neural growth through an alpha 7 nicotinic acetylcholine receptor mTOR pathway

**DOI:** 10.1101/2023.04.07.536081

**Authors:** Alexandru Graur, Nadine Kabbani

## Abstract

Acetylcholinesterase (AChE) is a highly conserved enzyme responsible for the regulation of acetylcholine signaling within the brain and periphery. AChE has also been shown to participate in non-enzymatic activity and contributing to development and aging. In particular, enzymatic cleavage of the carboxy terminal region of the synaptic AChE isoform, AChE-T, is shown to generate a bioactive T30 peptide that binds to the α7 nicotinic acetylcholine receptor (nAChR) at synapses. Here, we explore intracellular mechanisms of T30 signaling within the human cholinergic neural cell line SH-SY5Y using high performance liquid chromatography (HPLC) coupled to electrospray ionization mass spectrometry (ESI-MS/MS). Proteomic analysis of cells exposed to (100nM) T30 for 3-days reveals significant changes within proteins important for cell growth. Specifically, bioinformatic analysis identifies proteins that converge onto the mammalian target of rapamycin (mTOR) pathway signaling. Functional experiments confirm that T30 regulates neural cell growth via mTOR signaling and α7 nAChR activation. In addition, T30 was found promote mTORC1 pro-growth signaling through an increase in phosphorylated elF4E, and a decrease in autophagy LC3B-II level. Taken together, our findings define mTOR as a novel pathway activated by the T30 cleavage peptide of AChE and suggest a role for mTOR signaling in cholinergic aspects of brain development, as well as disease.

## Introduction

Acetylcholine (ACh) is an abundant neurotransmitter in the brain and periphery important for various physiological functions including movement, memory, and immune system regulation ^1^. The cholinergic synapse is among the most well understood synapses within many organisms, serving as a prototype for classical neurotransmission^2,3^. Amongst the primary molecular components of the cholinergic synapse are ACh binding receptors such as the ligand-gated nicotinic acetylcholine receptor channel (nAChR)^4^. In addition to their post-synaptic localization, nAChRs are also found presynaptically on cholinergic terminals and can contribute to synaptic growth and neurotransmitter release in brain circuits for memory and cognitive processing^5^. The α7 nAChR is a widespread homopentameric channel receptor that activates calcium within cells ^6^. Studies show that α7 nAChRs can signal through both ionotropic and metabotropic modes in neural and immune cells^7^. In particular, α7 nAChR signaling is important for neural cell development and synaptic growth ^8–10^.

The cholinergic synapse is marked by the presence of acetylcholinesterase (AChE), a powerful enzyme that regulates ACh levels within the synaptic cleft ^11^. AChE however is now well-established as a signaling molecule with dual hydrolytic and non-hydrolytic functions including strong trophic activity ^12–14^. The mammalian AChE gene contains six exons which are spliced in several alternative forms that create three main AChE isoforms (AChE-T,-R, -H) ^11,14^. The synaptic tetrameric variant AChE-T is the dominant isoform in the brain ^14^. AChE-T is a cell membrane attached enzyme via its well characterized proline-rich membrane anchor (PRiMA) domain ^11,11,14^. AChE-T also has an amphiphilic region within its c-terminus that contributes to oligomerization ^15^. Proteolytic cleavage of the last 30 amino acids at the c-terminus generates *in vitro*, as well as *in vivo* a T30 bioactive peptide ^16,17^. Interestingly, since the c-terminal region of AChE-T contains some sequence homology with the amyloid precursor protein (APP), cleavage of AChE-T and APP appears to yield two peptides (T30 and Aβ42, respectively) with some sequence similarity (**Fig. 1A)** and neurotoxic potential ^17^.

**Figure 1.**
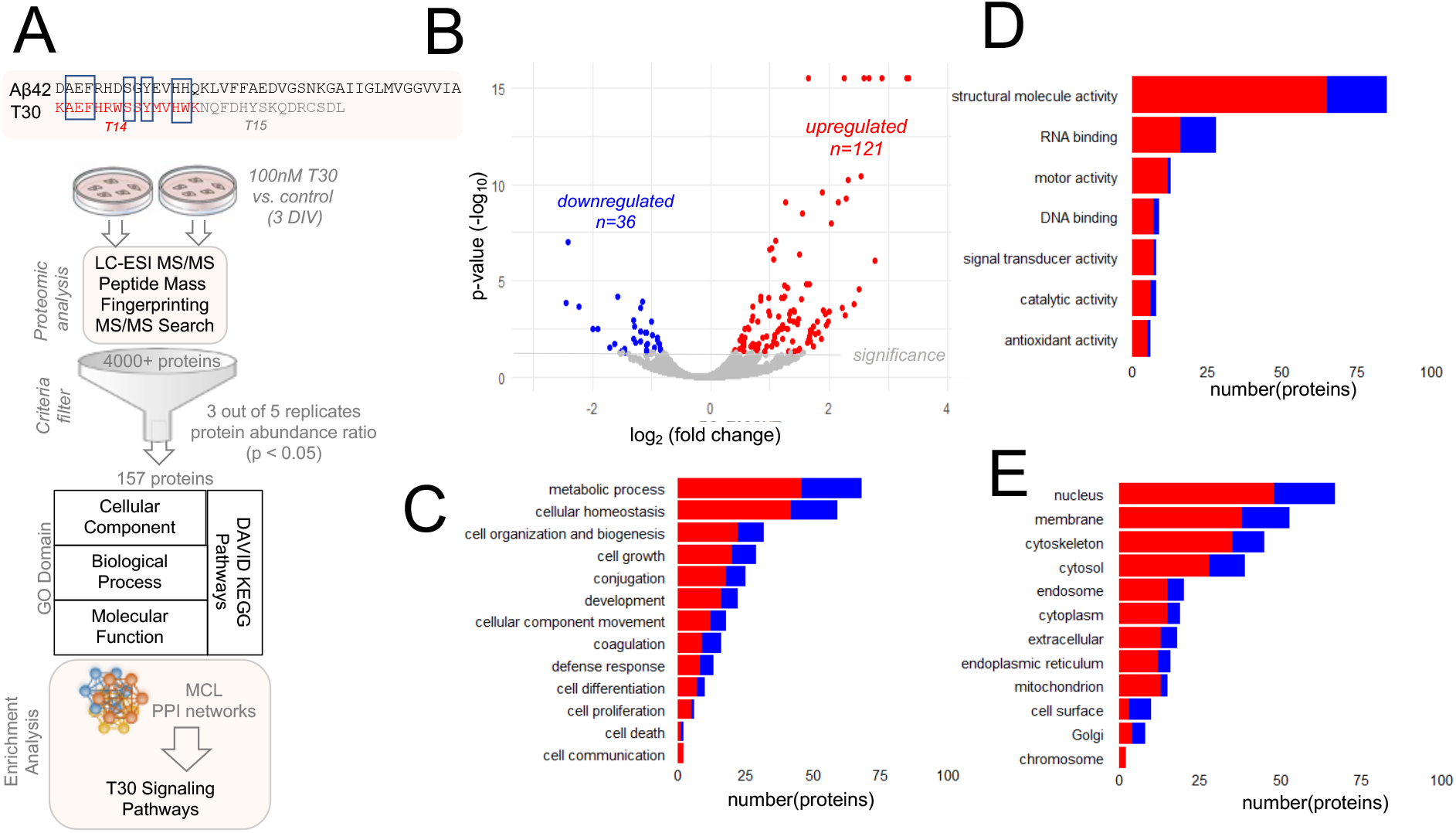
Proteomic analysis of T30 signaling. A) The T30 peptide sequence contains active (T14) and inactive (T15) portions as well as some homology to Aβ42. Cells were treated with 100nM T30 for 3 DIV then analyzed by LC-ESI MS/MS and bioinformatics. B) The distribution of detected proteins within T30 treated cells. The threshold for statistical significance (p < 0.05). C-E) Gene Ontology (GO) terms associated with significantly altered proteins (C: Biological Process, D: Molecular Function. E: Cellular Component)

Interestingly Aβ42 and T30 are reported to both bind to the α7 nAChR, and can thus impact calcium signaling and neurotransmission ^18–21^. The pharmacological targeting of T30 is a promising approach for neurodegenerative disease therapy ^22^. In this study we explore how T30 impacts signaling within the human neural cholinergic cell line SH-SY5Y, which is a model for Alzheimer’s Disease (AD) and known to endogenously express α7 nAChRs ^23,24^. Using quantitative whole cell proteomics and targeted functional cell assays, we identify a novel mechanism of T30 signaling via mTOR that leads to increased neural cell growth. T30 activation of mTOR may provide a mechanistic understanding of how non-enzymatic AChE-T can contribute to synaptic function.

## Methods

### Cell culture, transfection, and treatment

Human neuroblastoma cells SH-SY5Y cells (ATCC® CRL-2266™) were cultured in T75 flasks for propagation and then plated onto 100 μg/ml polyD-lysine (Millipore, A-003-E) coated 96-well glass bottom plates for imaging. Cells were fed DMEM (Gibco 11995065) supplemented with 10% fetal bovine serum (FBS) and 1% pen/strep at 37°C and 5% CO2. Experiments were conducted in cells that did not exceed 19 passages. For treatment experiments, cells were grown to 70% confluence then treated with: 100 nM T30/T15/ NBP14, 50 nM α-bungarotoxin (bgtx) (Thermo Fisher B1601), 1 μM Rapamycin (Thermo Scientific AAJ62473MF). Treatment media was changed daily. T30, T15, and NBP14 peptides were generously provided by NeuroBio LTD (Oxford, UK) and have been characterized elsewhere ^20,25,26^.

Cells were transfected using Lipofectamine 2000 (Thermo Fisher 11668030) with constructs encoding α7_345–348A_ in pEYFP-C1 ^27^ with the pEYFP-C1 plasmid used as a control. All cDNA constructs were propagated in DH5α cells (Thermo Fisher 18258012) and purified using a maxi prep kit (Xymo Research, Irvine, CA, USA). Cell proliferation was analyzed through live cell counting using phase contrast microscopy within a C-Chip hemocytometer (Bulldog Bio, Portsmouth, NH, USA).

### Protein Extraction and Western Blot

Proteins were obtained from cultured cells as previously described ^28^. In brief, at 3 DIV cells were lysed using a 0.1% Triton X-100 lysis buffer (Triton X-100, 150 mM NaCl, 20 mM Tris HCl, 2 mM EDTA, and 10% glycerol) supplemented with protease (Complete Mini, Roche) and PhosSTOP (Sigma Aldrich 4906845001) inhibitors. Protein concentration was determined using the Bradford assay. Proteins were separated on a NuPAGE 4-12% Bis-Tris gradient gel (Thermo Fisher NP0322BOX) and then transferred onto a nitrocellulose membrane (ThermoFisher IB301002). Membranes were blocked with milk prior to application of a primary antibody: GAPDH (1:1000 Cell Signaling 5174), LC3B (1:1000 Cell Signaling 2775), p-eIF4E (1:1000 Cell Signaling 9741), eIF4E (1:1000 Santa Cruz sc-271480) and Cytochrome C (1:1000 AbCam ab90529). HRP secondary antibodies were purchased from Jackson Immunoresearch (West Grove PA, USA). A SeeBlue Plus2 Ladder (ThermoFisher LC5925) was used as molecular weight marker. Bands were visualized using SuperSignal West Pico or SuperSignal West Femto Chemiluminescent substrates (ThermoFisher) via the G:BOX Imaging System and GeneSYS software (Syngene, Fredrick MD, USA). Band density was analyzed in Image J (NIH, Bethesda MD, USA). All measures were normalized to GAPDH unless otherwise stated. Average band intensity measures are based on three separate experiments (n = 3).

### Liquid-chromatography electrospray ionization mass spectrometry

Whole cell proteomic analysis was performed based on an established method ^19,29^. Briefly, solubilized protein samples were incubated for 5 minutes with acetone on ice followed by protein precipitation via centrifugation. The resulting protein pellet was denatured, reduced, and alkylated with 8 M urea, 1 M dithiothreitol, and 0.5 M iodoacetamide. Proteins were digested with trypsin (0.5 μg/μl) in 500nM ammonium bicarbonate and incubated at 37°C for 5 hrs. The samples were then desalted with C-18 ZipTips (Millipore), dehydrated in a SpeedVac for 18 mins and reconstituted in 0.1% formic acid before undergoing liquid-chromatography electrospray ionization mass spectrometry (LC-ESI MS/MS) with 5 technical replicates.

LC-ESI MS/MS was performed using an Exploris Orbitrap 480 equipped with an EASY-nLC 1200HPLC system (Thermo Fischer Scientific, Waltham, MA, USA). Peptides were separated using a reverse-phase PepMap RSLC 75 μm i.d by 15 cm long with a 2 μm particle size C18 LC column (Thermo Fisher Scientific, Waltham, MA, USA), and eluted with 80% acetonitrile and 0.1% formic acid at a flow rate of 300 nl/min. After a full scan at 60,000 resolving power from 300 m/z to 1200 m/z, peptides were fragmented by high-energy collision dissociation (HCD) with a normalized collision energy of 28%. EASY-IC filters for monoisotopic precursor selection, internal mass calibration, and dynamic exclusions (20 s) were enabled. Data on peptide precursor ions with charge states from +2 to + 4 was recorded.

### Proteomic quantification and statistical analysis

The SEQUEST HT search engine within the Proteome Discoverer *v*2.4 (Thermo Fisher Scientific, Waltham, MA, USA) was used to identify proteins by comparing raw MS peptide spectra to the NCBI 2018 human protein database using the following search engine parameters: mass tolerance for precursor ions = 2 ppm; mass tolerance for fragment ions = 0.05 Da; and cut-off value for the false discovery rate (FDR) in reporting peptide spectrum matches (PSM) to the database = 1%. Peptide abundance ratios were determined by precursor ion quantification in Proteome Discoverer *v*2.4, with the vehicle control group used as the denominator. Statistically significant abundance ratios with adjusted p-values < 0.05 were determined using a one-way analysis of variance (ANOVA) followed by Benjamini-Hochberg post-hoc analyses. Proteins with a quantifiable spectra signal profile seen in at least 3 of the 5 technical replicates were included in the analysis. Markov Cluster Algorithm (MCL) with an inflation parameter of 3 was used to perform clustering analysis on the data in the STRING database. Data was analyzed, organized, and presented using the R package (R Core Team, 2021): ggplot2 ^30^, tidyverse ^31^, Excel, the Database for Annotation, Visualization, and Integrated Discovery (DAVID) and Search Tool for the Retrieval of Interacting Genes/Proteins (STRING, *v*11.5) application ^32^.

### Immunocytochemistry and cell imaging

SH-SY5Y cells were fixed in a solution consisting of 1x PEM (80 mM PIPES, 5 mM EGTA, and 1mM MgCl2, pH 6.8) and 0.3% glutaraldehyde then quenched with sodium borohydride (2 mg/ml). Cells were permeabilized using 0.05% Triton X-100 (Sigma Aldrich). Quantification of structural change (i.e., neurite shape and growth) was performed using f-actin labeling with rhodamine phalloidin (Cytoskeleton PHDG1-A). All morphometric measures were conducted and quantified using ImageJ (NIH, Bethesda, MD, USA) as described ^8^. α7 nAChRs were detected at the cell surface and within the cytoplasm using Alexa Fluor 488 conjugated α-bungarotoxin (Alexa-488 bgtx) (Thermo Fisher B13422) as described ^27^. Images were captured using an inverted Zeiss LSM800 confocal microscope and the Zen software package (Carl Zeiss AG, Oberkochen, Germany).

## Results

### Identification of a T30 reactive proteome within neural cells

The human neuroblastoma SH-SY5Y cell line is a well-established model for the study of neural cell development and neurodegeneration ^24,33^. SH-SY5Y cells maintain the ability to model cholinergic neurons with endogenous expression of various cholinergic receptors ^23,34^. We used SH-SY5Y cells to examine proteomic responses to treatment with the bioactive c-terminal peptide of the AChE-T enzyme T30 ^25^. Cells were treated with 100nM T30 for 3 DIV then processed for proteomic analysis. T30 has been identified as an endogenous ligand of the α7 nAChR, and at 100nM it is shown to activate α7 nAChR calcium signaling in various cell lines ^25,35^. In these experiments, we used the vehicle (water) treatment condition as the control group in protein comparison.

We have developed a shotgun LC/ESI-MS/MS approach to identify cellular proteins, for SH-SY5Y and other cell lines, in response to various stimuli ^19,29^. In this study, a similar LC/ESI-MS/MS peptide detection method was used and quantification of protein changes was conducted based on label-free precursor ion abundance ratio measures between T30 treated cells and control samples (**Fig. 1A)**. Our proteomic analysis identified 4331 cellular proteins within the sample. A volcano plot distribution (**Fig. 1B**) shows that 121 of these proteins were significantly increased while 40 were significantly decreased and 4170 did not statistically change between the two experimental conditions (p < 0.05). Using Proteome Discoverer, we annotated significantly altered proteins according to the three main GO domains: biological processes, molecular function, and cellular components. GO domain terms matching the greatest number of altered proteins in response to T30 treatment are presented in **Fig. 1C-E**. A full list of the significantly altered proteins and peptide scores within the T30 treatment condition is provided **Supplemental Table 1**.

Whole cell proteomics enables an analysis of functional changes within cells that can be gleaned from examining protein-protein interaction (PPI) networks ^36^. Modifications to PPI networks can reveal important information on functional adaptive responses to an extracellular signal, as was done in Kabbani, 2008. We used a Markov cluster (MCL) analysis to define PPI networks within the T30 proteome ^38^. MCL analysis shows a relatively integrated PPI network based on the identity of the significantly altered proteins within SH-SY5Y cells (**Fig. 2**). Within this PPI network we identified various significantly altered protein clusters and functional pathways. The largest PPI cluster was found to contain 19 proteins with 53 connections yielding a significant PPI enrichment (p < 1.11 × 10^−16^). This cluster (Cluster 1) consisted of an average local clustering coefficient (ALCC) of 0.697. Enrichment analysis of Cluster 1 confirms ribosome enrichment in KEGG Pathways with a false discovery rate (FDR) of 8.49 × 10^−7^. MCL analysis revealed 11 clusters within the T30-associated PPI network (**Fig. 2 and Table 1)**. Many of the identified clusters were involved in pathways for cell growth including protein regulation.

**Figure 2.**
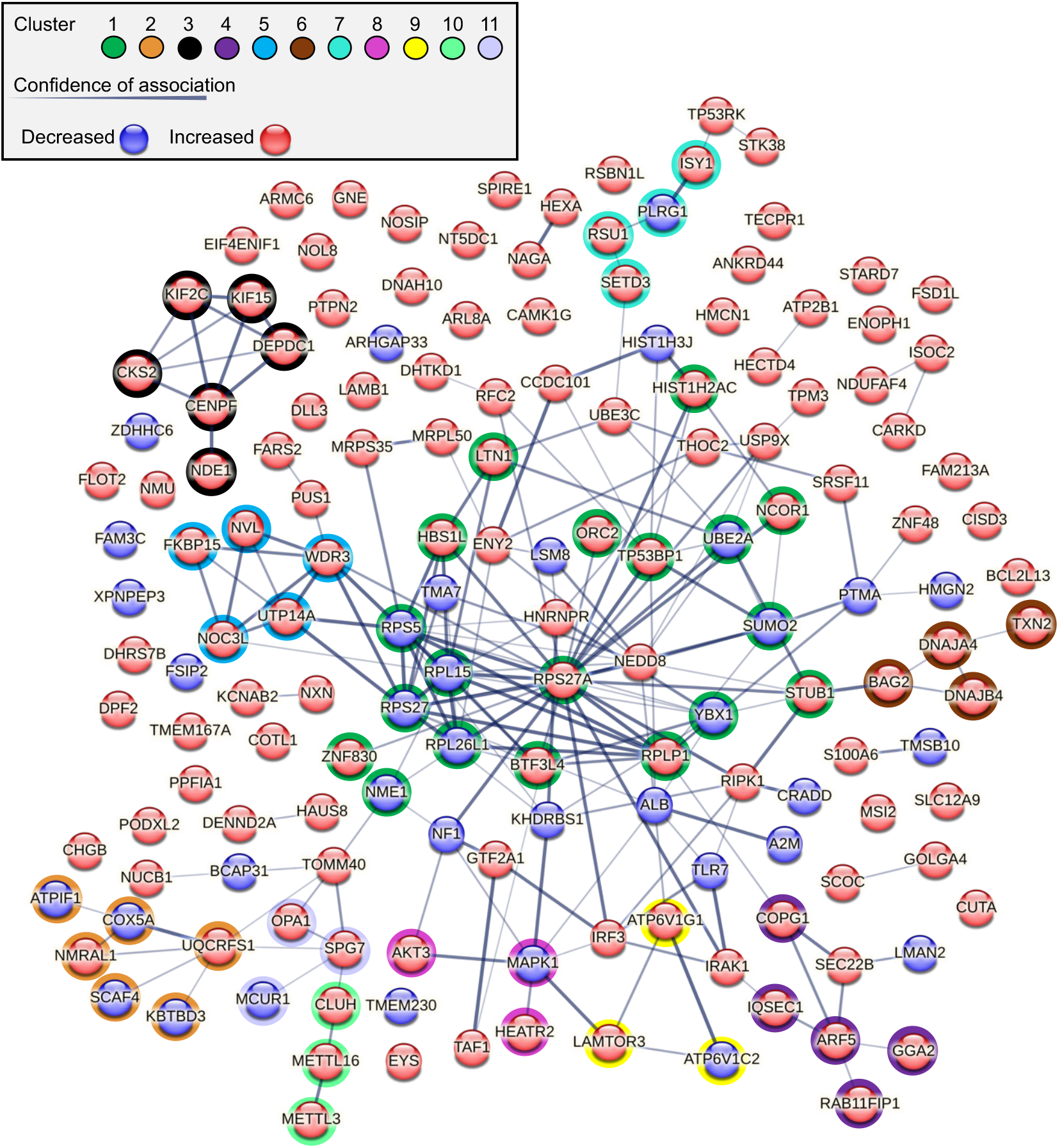
STRING network and cluster analysis of the T30 proteome. STRING analysis of the T30 proteome (all significantly altered proteins) showing network networks for interacting proteins. Line thickness reflects the confidence between node associations, and node color indicates whether the protein is increased or decreased. A Markov cluster algorithm (MCL) was used to identify 11 functional clusters within the proteome network.

**Table 1.** Top clusters identified using MCL in STRING

| Cluster # | Node # | Edge # | ALCC | PPIE | Cluster Enrichment |
| --- | --- | --- | --- | --- | --- |
| 1 | 19 | 53 | 0.697 | 1.11E-16 | Ribosome |
| 2 | 6 | 6 | 0.75 | 6.95E-09 | Electron transport chain/ Mitochondria |
| 3 | 6 | 11 | 0.933 | 5.99E-13 | Microtubule binding |
| 4 | 5 | 4 | 0.8 | 2.71E-07 | Protein transport |
| 5 | 5 | 9 | 0.9 | 5.71E-10 | Ribosome biogenesis in eukaryotes |
| 6 | 4 | 4 | 0.833 | 1.26E-07 | Chaperone |
| 7 | 4 | 3 | 0.5 | 1.78E-05 | Spliceosome |
| 8 | 3 | 2 | 0.667 | 8.47E-03 | mTOR signaling pathway |
| 9 | 3 | 3 | 1 | 1.16E-06 | MTORC1 regulation |
| 10 | 3 | 2 | 0.667 | 4.33E-05 | RNA binding |
| 11 | 3 | 2 | 0.667 | 8.58E-05 | Mitochondrial calcium ion transmembrane transport |

### T30 activates an mTOR pathway for cell growth

Studies have shown a role for non-hydrolytic AChE function in neuronal growth and synaptic maturation ^13,39^. In particular, the c-terminal fragment (T30) produced by AChE-T cleavage has been shown to activate intracellular signaling important for neural cell development ^40,41^. Our proteomic analysis reveals new proteins and pathways that are altered in response to a 3-day T30 presentation within the SH-SY5Y cell line. Bioinformatic KEGG Pathway analysis in DAVID further revealed enrichment of proteins involved in mTOR pathway signaling. Further analysis indicates that mTOR can serve as a point of convergence between PPI networks and several of the identified clusters within dataset (**Fig. 3A**). In **Fig. 3B**, a mechanistic pathway summarizing the hypothesized involvement of mTOR pathway in T30 mediated growth is presented. In this pathway, differentially altered (increased and decreased) proteins identified within the proteome are found as important components of mTOR signaling. The proteomic data suggests that T30 presentation promotes mTORC1 through the regulation of downstream signaling proteins.

**Figure 3.**
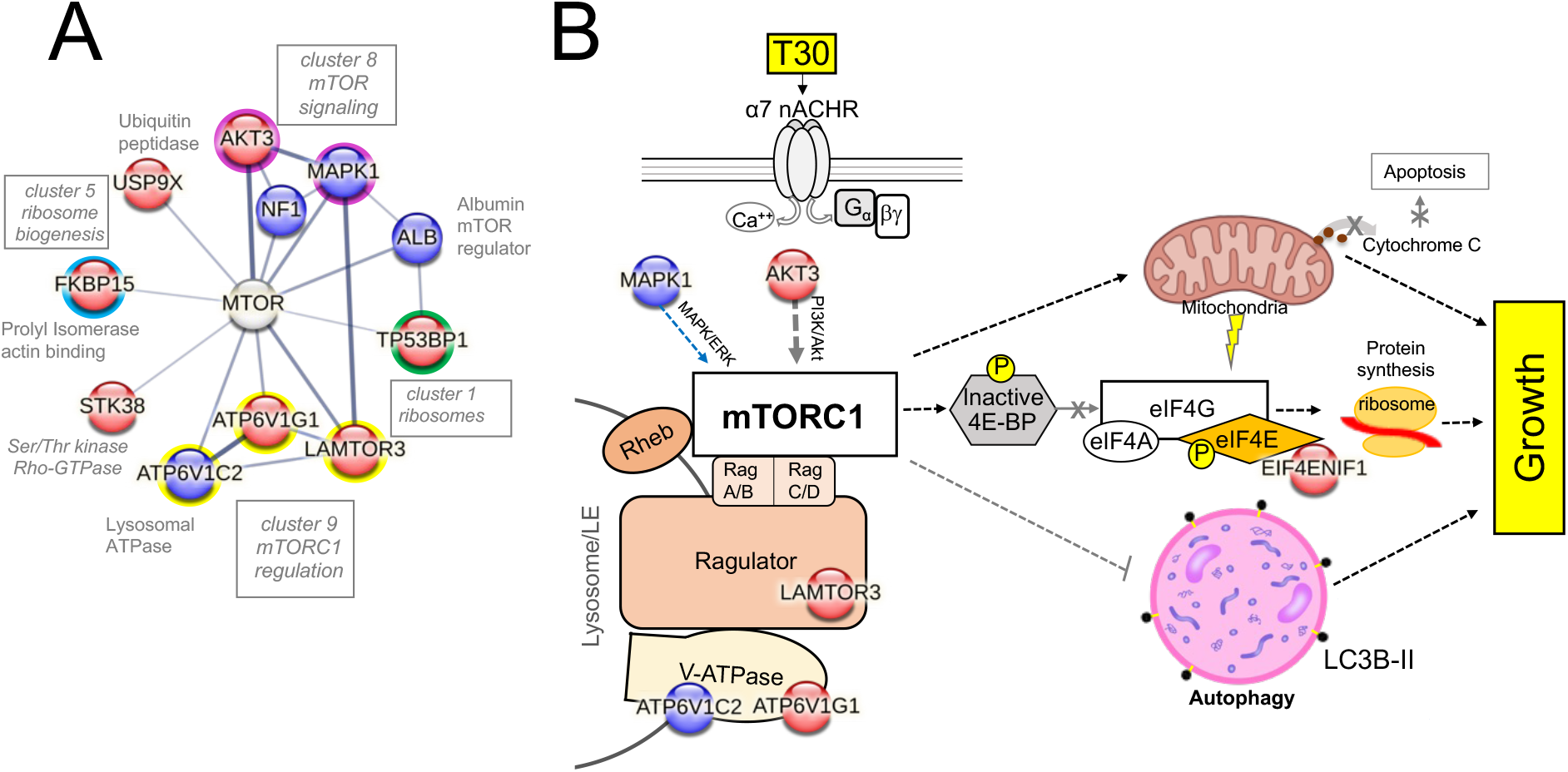
Identification of the mTOR pathway as a cellular target of T30 action. A) mTOR is found at the convergence for several proteins and MCL clusters (1, 5, 7, 9). B) A signal transduction hypothesis model for T30 activation of mTOR in cells. Proteins discovered through the proteomic assay are indicated within the red (increased) and blue (decreased) circles.

To test this, we examined SH-SY5Y cell growth in the presence of the 100nM T30 for 3 DIV. As shown in **Fig. 4A**, T30 treatment increased cell proliferation but this effect was not found to be statistically significant (p=0.287). Protruding from the membranes of developing neural cells are motile structures that consist of actin projecting lamellipodia as well as cytoplasmic filopodia ^42,43^. In previous studies we have shown a role for α7 nAChRs in regulating actin-mediated cytoskeletal growth in neurites and growth cones ^8,28^. T30 presentation showed a significant increase in neurite growth as measured by neurite number, increased presence of filipodia/lamellipodia structures, and total measured surface area (n=40, p<.001). This effect was not seen in response to application of the cyclic variant of the T30 peptide NBP14 shown to be biologically inactive (Brai et al., 2018) (n=40, p=0.369) (**Fig. 4B-C**).

**Figure 4.**
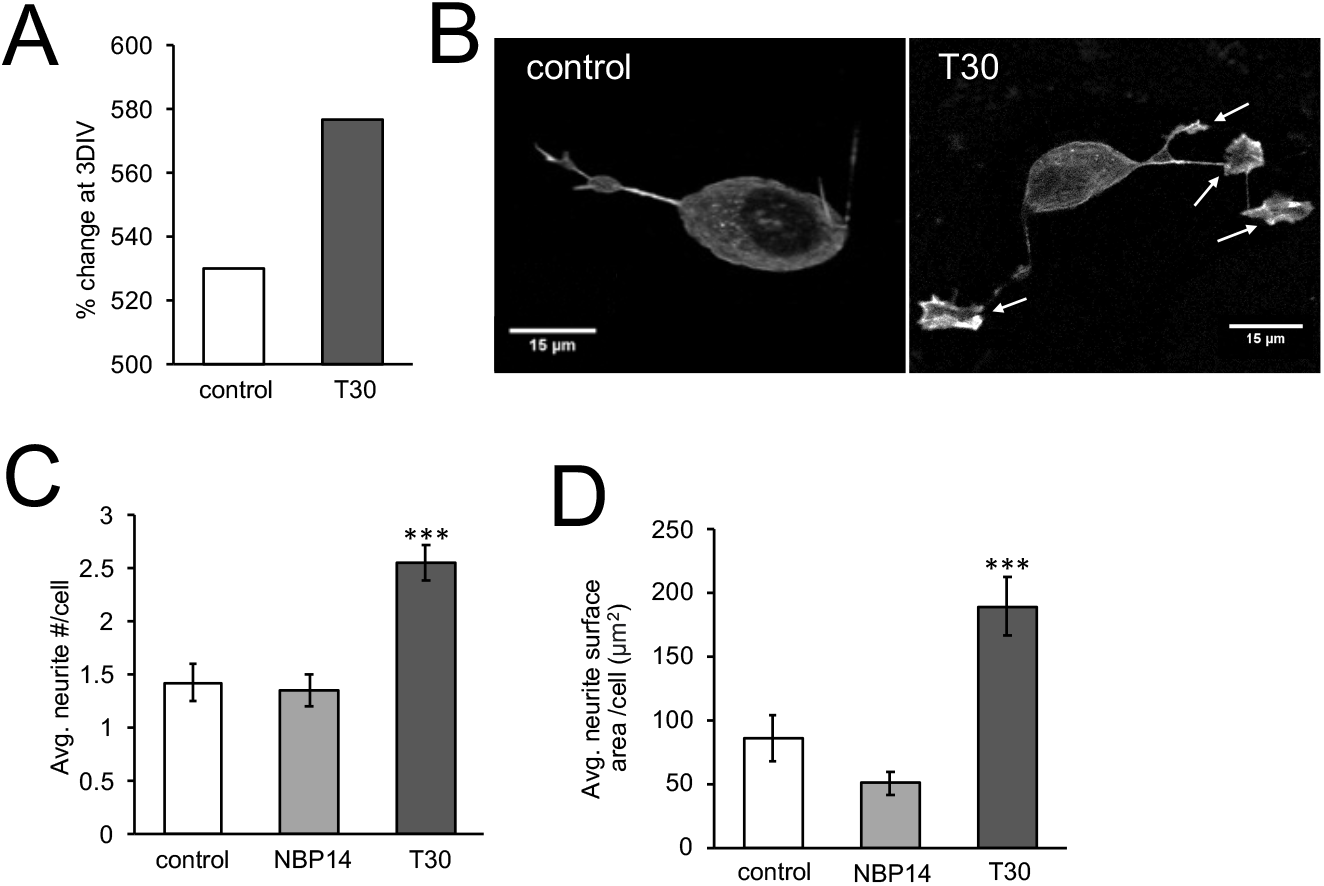
T30 treatment regulates neural growth. SH-SY5Y cells were treated with 100nM T30 for 3 days in vitro (DIV) then imaged using fluorescent (f-actin) phalloidin. A) Cell proliferation shown as a percentage change in total cell number after 3 DIV. B) Representative cell images at 3 DIV. Arrows point to growth filipodia and lamellipodia structures. C) Average neurite number per cell. D) Average neurite surface area per cell. n=40,*** p<0.001

### A requirement for α7 nAChR signaling in T30-mediated growth

Studies show that T30 binds to α7 nAChRs in neural cells activating intracellular calcium signaling and increasing nAChR expression ^44^. We confirmed the role of α7 nAChRs in T30-mediated neurite growth using the selective α7 nAChR blocker α-bungarotoxin (bgtx). As shown in **Fig. 5A**, co-treatment of cells with 50nM bgtx and T30 did not produce an effect on growth (n=40, p=0.136). We examined the impact of T30 treatment on the expression and localization of the α7 nAChR within SH-SY5Y cells. Fluorescence imaging was performed using Alexa 488-bgtx to assess α7 nAChR expression as previously shown ^27^. We first compared cell surface and intracellular nAChR expression by labeling cells with Alexa 488-bgtx under non-permeabilized and permeabilized fixation conditions, respectively. Data shows that T30 treatment increases the Alexa 488-bgtx signal within the cell (n=40, p <0.05) but not at the cell surface (n=40, p=0.139) (**Fig. 5B)**. We next examined the effect of T30 on α7 nAChR expression in growth sites. Our previous findings show that this nAChR is targeted to growth cones and can directly regulate structural motility through cytoskeletal modulation ^8,45^. As shown in **Fig. 5C**, treatment with T30 was found to significantly increase the Alexa 488-bgtx signal at terminal growth sites relative to both the control condition (n=30, p=0.0002) as well as the cyclic NBP14 peptide (n=30, p=0.005).

**Figure 5.**
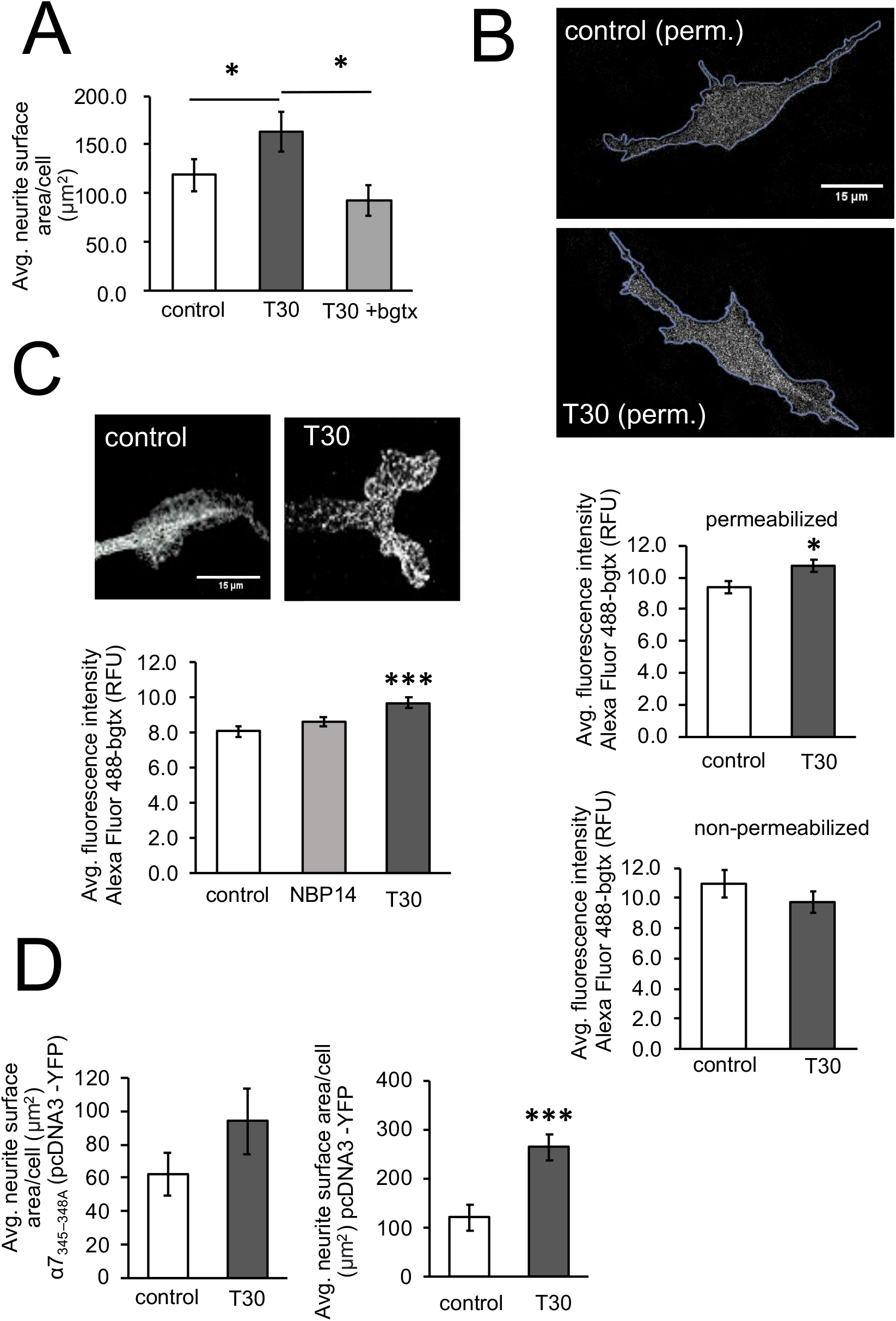
T30 activates α7 nAChR signaling to promote growth. A) Average neurite surface area at 3 DIV. B) Images, representative permeabilized cell images showing ROI analysis of Alexa-488-bgtx signal intensity. Histograms, average fluorescence intensity of the Alexa-488-bgtx signal in permeabilized and non-permeabilized cells. C) Average fluorescence intensity of the Alexa-488-bgtx signal within sites of growth. D) Average neurite surface area measures within cells transfected with α7_345–348A_ or pEYFP-C1. n=40, * p<0.05, *** p<0.001

The activation of the α7 nAChR can regulate axonal development within hippocampal neurons through the ability of the α7 nAChR to directly bind and activate heterotrimeric GTP-binding proteins (G proteins) ^27^. Expression of a mutant α7 subunit (α7_345–348A_) that lacks the G protein-binding site has been established as a method for blocking α7 nAChR-mediated G protein signaling and growth ^27^. We tested the ability of T30 to promote growth in SH-SY5Y cells transfected with α7_345–348A_ (pcDNA3-YFP). In this assay, control cells were transfected with the expression vector pcDNA3-YFP alone. Morphological analysis shows that T30 treatment does not increase neurite growth in cells expressing α7_345–348A_ (n=40, p=0.09) **(Fig. 5D)**.

### T30 activates an mTOR signaling pathway

Our proteomic analysis reveals an enrichment of intracellular proteins involved in mTOR signaling (**Fig. 3 and Table 1**). Activation of the mTOR pathway is a conserved evolutionary signaling strategy for balancing cell growth and managing metabolic demands in various environmental contexts ^46–48^. Previous work indicates that α7 nAChRs activate mTOR during cellular development, inflammation, and cancer cell progression ^49^. We directly confirmed the involvement of mTOR signaling in T30-mediated neurite growth using the mTOR inhibitor rapamycin. As shown in **Fig. 6A**, pre-treatment of neural cells with 1uM rapamycin, for 24 hours, was found sufficient to abolish the effect of T30 on neurite growth (n=40, p=0.161). The activation of mTORC1 is shown to promote the phosphorylation of the translation regulating factor eIF4E (serine 209) as a key component of mTOR-mediated translational regulation and growth (Kosciuczuk et al., 2017; Majeed et al., 2021, **Fig. 3B**). We assessed the effect of T30 treatment on eIF4E expression and phosphorylation at 3 DIV. As shown in **Fig. 6B**, T30 treatment did not increase eIF4E expression but significantly increased its phosphorylation (phospho-eIF4E (p=0.011)).

**Figure 6.**
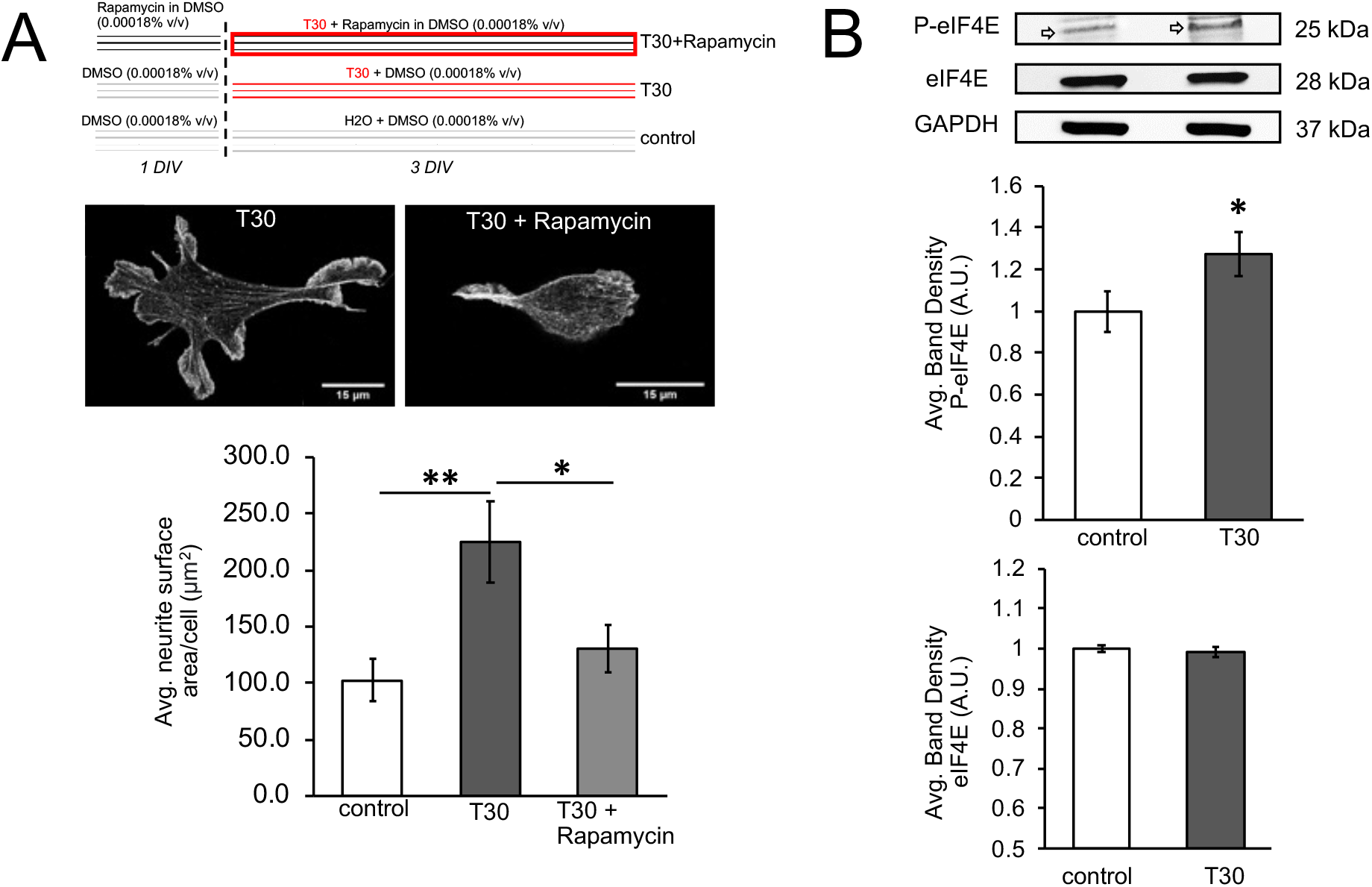
Analysis of the mTOR pathway in T30 mediated growth. A) Top, summary of the rapamycin experiment. Middle images, representative cells at 3 DIV. Bottom histogram, average neurite surface area measures. B) Top, representative immunoblots. Bottom, average band density measures from 3 separate experiments. n=40, * p<0.05, ** p<0.005

The mTOR pathway plays a critical role in maintaining cellular balance between anabolic and catabolic states through the regulation of degradation-mediated autophagy processes ^52^. Isoforms of the cytosolic light chain (LC3) protein undergo modifications during autophagy and thus serve as important autophagy markers ^53^. We examined autophagy-mediated LC3I to II conversion within LC3B as previously shown ^53^. LC3B was detected throughout the cell, including sites of growth, consistent with the role of the autophagosome in modulating neural growth ^54^ (**Fig. 7A**). Treatment of cells with T30 (for 3 DIV) was found to significantly reduce LC3B-II levels consistent with mTORC1-mediated autophagy inhibition (**Fig. 7B**). The ability of T30 to reduce LC3B-II was blocked by co-application of bgtx (p=0.428) consistent with the role of the α7 nAChR in T30 effect autophagy regulation.

**Figure 7.**
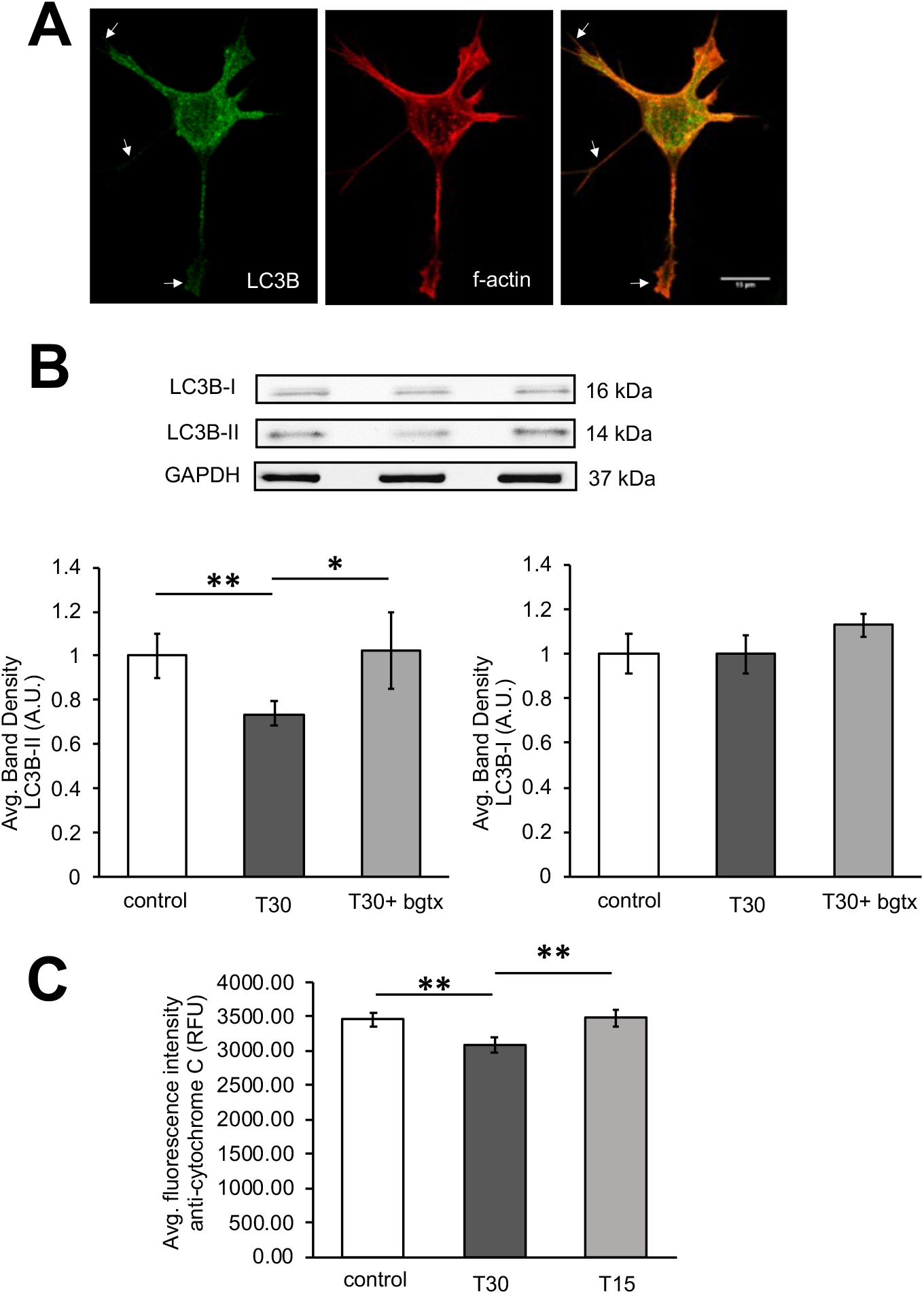
T30 treatment is associated with a reduction in autophagy and cytochrome c. A) Representative cell images at T30 treatment at 3DIV. Arrows point to localization of the LC3B autophagy marker within sites of growth. B) Top, representative immunoblots. Bottom, average band density measures. C) Average fluorescence intensity measures of the anti-cytochrome c immunosignal. n=40, * p<0.05, ** p<0.005

In mammalian cells, the mTOR pathway coordinates mitochondrial energy production and stimulates the synthesis of various mitochondrial proteins ^55^. Our proteomic analysis indicates an effect of T30 on the regulation of mitochondrial proteins (**Fig. 2 and Table 1**). Indeed, AChE and α7 nAChR are individually documented to regulate mitochondrial activity and to contribute to apoptotic signaling in neurons ^56,57^. We examined the effect of the T30 treatment on cytochrome C levels within SH-SY5Y cells. Immunofluorescence analysis using an anti-cytochrome C antibody shows that T30 treatment reduces cytochrome C within cells relative to vehicle treated controls (n=40, p=0.009). In these experiments, the non-bioactive portion of T30 (T15) did not have an effect on cytochrome C in cells (**Fig. 7C**) (n=40, p=0.459).

## Discussion

AChE is an enzyme vital for mammalian synaptic transmission through its ability to hydrolyze ACh ^58,59^. It is also widely expressed outside of the nervous system and is sometimes found in non-cholinergic cells ^60^. A large body of work has demonstrated non-hydrolytic functional properties for AChE within various cell types ^12,13,61^. Amongst its non-hydrolytic activity is its role as a trophic factor in various cell types including bone and cancer ^62^. Studies show that synaptic AChE-T is especially abundant during brain development and can regulate axonal growth as well as pathfinding during synaptogenesis ^63^. The loss of AChE-T in mice is associated with disruption to synaptic connectivity within the retina and cortex ^64^. Our study supports the involvement of non-hydrolytic AChE-T in growth demonstrating an important role for the T30 peptide in human neural cells ^65,66^.

Earlier findings demonstrate that T30 acts by directly binding and activating signaling through the α7 nAChR ^44^. Sequence similarity between T30 and Aβ42 has been shown yet it is not yet clear if the two peptides share common binding and function at the nAChR. Interestingly, in *ex vivo* rat brain slices, the application of T30 results in an increase in the expression of Aβ42 ^67^, suggesting that T14 can interfere with amyloid protein turnover amd management. Interactions between AChE and nAChR have also been explored in various context ^68^. Thus, while the two molecules are co-expressed at the mature cholinergic synapse, AChE-T and the α7 nAChR appear highly coupled in expression during early brain development^69^. During early post-natal synaptic development, α7 nAChR is at its highest within rodent brain and shown to regulate neural cell proliferation and synaptic maturation within brain regions such as the hippocampus ^10^. Our earlier studies have shown an important role for α7 nAChR metabotropic signaling through G proteins in axonal growth and calcium signaling at the growth cone ^28^. In this study, T30 activation of the α7 nAChR is also able to support neurite growth through a process dependent on G protein signaling since the expression of the α7_345–348A_ mutant blocked the ability of T30 to mediate growth.

Our proteomic analysis reveals several important intracellular pathways that appear engaged by the presentation of T30 *in vitro*. These pathways all appear to promote cell growth and support protein synthesis. In fact, even when looking at all statistically altered proteins within the T30-associated proteome, ∼75% of the change was due to an increase in the expression of specific proteins. Bioinformatic analysis using MCL and DAVID KEGG pathway indicates that these protein changes reflect an mTOR pro-growth state within T30 treated cells. Indeed, mTOR is an evolutionarily conserved serine/threonine kinase that regulates many cellular responses (from autophagy to translation) and responds to incoming signals by modulating energy availability and metabolic demand ^70,71^. It functions via two distinct complexes: mTORC1 and mTORC2 with the activation mTORC1 resulting in protein synthesis via p70 S6 kinase (S6K1 and S6K2) and phosphorylation of eukaryotic initiation factor 4-binding protein (4EBP1 and 4EBP2). mTORC1 also suppresses autophagy mediated protein degradation and can thus contribute to growth ^52^. Our findings show that T30 promotes neural cell growth by activating α7 nAChRs and mTOR pathway signaling. This process is supported by recently published evidence on the ability of α7 nAChRs to support AKT/mTOR autophagy within neurons^72,73^.

Our experiments show that T30 acts via the mTOR pathway at several points, first by decreasing the expression of the autophagy marker LC3B and second by increasing the phosphorylation of eIF4E. Both of these processes are likely to promote neurite growth and explain the actions of T30 on growth within our cells as well as elsewhere ^50,52^. Interestingly, the effects of T30 appear accompanied by overall reduction in cellular cytochrome c, which is a driver of apoptosis ^74^. The effects of T30 are found specific since antagonism of the α7 nAChR with bgtx abolished T30-associated growth, and application of the mTOR inhibitor rapamycin and non-bioactive peptide variants of T30 (T15 and NBP14) did not promote in signaling nor growth.

Disruption to mTOR signaling is implicated in many human disease including auto-immune disorder, neurodegeneration, and various cancers ^46^. In the brain high stimulation of mTOR has been suggested to promote hyperphosphorylation of synaptic tau and drive amyloid protein accumulation ^75^. A growing body of evidence demonstrates a link between mTOR signaling and AD through protein. For example, an alteration in the autophagy-lysosome pathway has been shown to drive Aβ42 neurotoxicity ^76,77^, and a loss in mTORC1 regulation appears to contribute to protein aggregation within neural cells ^78^. It has been suggested that interactions between T30 and nAChRs can participate in early cholinergic cell death within the brain ^79^. This study provides novel evidence on a connection between the mTOR pathway and T30-mediated α7 nAChR signaling. How this may contribute to neural development and disease is an important question for the future with significant therapeutic implications.

## References

1. Haam, J. & Yakel, J. L. Cholinergic modulation of the hippocampal region and memory function. J. Neurochem. 142, 111–121 (2017).

2. Alkondon, M., Pereira, E. F., Cortes, W. S., Maelicke, A. & Albuquerque, E. X. Choline is a selective agonist of alpha7 nicotinic acetylcholine receptors in the rat brain neurons. Eur. J. Neurosci. 9, 2734–2742 (1997).

3. Albuquerque, E. X., Pereira, E. F. R., Alkondon, M. & Rogers, S. W. Mammalian nicotinic acetylcholine receptors: from structure to function. Physiol. Rev. 89, 73–120 (2009).

4. Cecchini, M. & Changeux, J.-P. The nicotinic acetylcholine receptor and its prokaryotic homologues: Structure, conformational transitions & allosteric modulation. Neuropharmacology 96, 137–149 (2015).

5. Lendvai, B., Kassai, F., Szájli, Á. & Némethy, Z. α7 Nicotinic acetylcholine receptors and their role in cognition. Brain Res. Bull. 93, 86–96 (2013).

6. Shen, J. & Yakel, J. L. Nicotinic acetylcholine receptor-mediated calcium signaling in the nervous system. Acta Pharmacol. Sin. 30, 673–680 (2009).

7. Kabbani, N. et al. Are nicotinic acetylcholine receptors coupled to G proteins? BioEssays 35, 1025–1034 (2013).

8. King, J. R. & Kabbani, N. Alpha 7 nicotinic receptor coupling to heterotrimeric G proteins modulates RhoA activation, cytoskeletal motility, and structural growth. J. Neurochem. 138, 532–545 (2016).

9. King, J. R. & Kabbani, N. Alpha 7 nicotinic receptors attenuate neurite development through calcium activation of calpain at the growth cone. PLOS ONE 13, e0197247 (2018).

10. Lozada, A. F. et al. Glutamatergic Synapse Formation is Promoted by α7-Containing Nicotinic Acetylcholine Receptors. J. Neurosci. 32, 7651–7661 (2012).

11. Richbart, S. D., Merritt, J. C., Nolan, N. A. & Dasgupta, P. Acetylcholinesterase and human cancers. in Advances in Cancer Research vol. 152 1–66 (Elsevier, 2021).

12. Halliday, A. C. & Greenfield, S. A. From protein to peptides: a spectrum of non-hydrolytic functions of acetylcholinesterase. Protein Pept. Lett. 19, 165–172 (2012).

13. Silman, I. & Sussman, J. L. Acetylcholinesterase: ‘classical’ and ‘non-classical’ functions and pharmacology. Curr. Opin. Pharmacol. 5, 293–302 (2005).

14. Zimmermann, M. Neuronal AChE splice variants and their non-hydrolytic functions: redefining a target of AChE inhibitors? Br. J. Pharmacol. 170, 953–967 (2013).

15. Heider, H. & Brodbeck, U. Monomerization of tetrameric bovine caudate nucleus acetylcholinesterase. Implications for hydrophobic assembly and membrane anchor attachment site. Biochem. J. 281 (Pt 1), 279–84 (1992).

16. Jean, L., Thomas, B., Tahiri-Alaoui, A., Shaw, M. & Vaux, D. J. Heterologous amyloid seeding: revisiting the role of acetylcholinesterase in Alzheimer’s disease. PloS One 2, e652 (2007).

17. Garcia-Ratés, S. & Greenfield, S. When a trophic process turns toxic: Alzheimer’s disease as an aberrant recapitulation of a developmental mechanism. Int. J. Biochem. Cell Biol. 149, 106260 (2022).

18. Dineley, K. T. Beta-amyloid peptide--nicotinic acetylcholine receptor interaction: the two faces of health and disease. Front. Biosci. J. Virtual Libr. 12, 5030–5038 (2007).

19. Sinclair, P. & Kabbani, N. Nicotinic receptor components of amyloid beta 42 proteome regulation in human neural cells. PLOS ONE 17, e0270479 (2022).

20. Garcia-Ratés, S. et al. (I) Pharmacological profiling of a novel modulator of the α7 nicotinic receptor: Blockade of a toxic acetylcholinesterase-derived peptide increased in Alzheimer brains. Neuropharmacology 105, 487–499 (2016).

21. Xu, C., Zhao, L. & Dong, C. A Review of Application of Aβ42/40 Ratio in Diagnosis and Prognosis of Alzheimer’s Disease. J. Alzheimers Dis. JAD 90, 495–512 (2022).

22. Greenfield, S. A. et al. A novel process driving Alzheimer’s disease validated in a mouse model: Therapeutic potential. Alzheimers Dement. Transl. Res. Clin. Interv. 8, e12274 (2022).

23. Elnagar, M. R. et al. Functional characterization of α7 nicotinic acetylcholine and NMDA receptor signaling in SH-SY5Y neuroblastoma cells in an ERK phosphorylation assay. Eur. J. Pharmacol. 826, 106–113 (2018).

24. Bell, M. & Zempel, H. SH-SY5Y-derived neurons: a human neuronal model system for investigating TAU sorting and neuronal subtype-specific TAU vulnerability. Rev. Neurosci. 33, 1–15 (2022).

25. Bond, C. E., Zimmermann, M. & Greenfield, S. A. Upregulation of α7 Nicotinic Receptors by Acetylcholinesterase C-Terminal Peptides. PLoS ONE 4, e4846 (2009).

26. Cottingham, M. G., Hollinshead, M. S. & Vaux, D. J. T. Amyloid Fibril Formation by a Synthetic Peptide from a Region of Human Acetylcholinesterase that Is Homologous to the Alzheimer’s Amyloid-β Peptide. Biochemistry 41, 13539–13547 (2002).

27. King, J. R., Nordman, J. C., Bridges, S. P., Lin, M.-K. & Kabbani, N. Identification and Characterization of a G Protein-binding Cluster in α7 Nicotinic Acetylcholine Receptors. J. Biol. Chem. 290, 20060–20070 (2015).

28. Nordman, J. C. & Kabbani, N. An interaction between α7 nicotinic receptors and a G-protein pathway complex regulates neurite growth in neural cells. J. Cell Sci. 125, 5502–5513 (2012).

29. Sinclair, P., Baranova, A. & Kabbani, N. Mitochondrial Disruption by Amyloid Beta 42 Identified by Proteomics and Pathway Mapping. Cells 10, 2380 (2021).

30. Wickham, H. ggplot2: Elegant graphics for data analysis. (2016).

31. Wickham, H. et al. Welcome to the Tidyverse. J. Open Source Softw. 4, 1686 (2019).

32. Szklarczyk, D. et al. The STRING database in 2021: customizable protein-protein networks, and functional characterization of user-uploaded gene/measurement sets. Nucleic Acids Res. 49, D605–D612 (2021).

33. Tagai, N., Tanaka, A., Sato, A., Uchiumi, F. & Tanuma, S.-I. Low Levels of Brain-Derived Neurotrophic Factor Trigger Self-aggregated Amyloid β-Induced Neuronal Cell Death in an Alzheimer’s Cell Model. Biol. Pharm. Bull. 43, 1073–1080 (2020).

34. Groot Kormelink, P. J. & Luyten, W. H. Cloning and sequence of full-length cDNAs encoding the human neuronal nicotinic acetylcholine receptor (nAChR) subunits beta3 and beta4 and expression of seven nAChR subunits in the human neuroblastoma cell line SH-SY5Y and/or IMR-32. FEBS Lett. 400, 309–314 (1997).

35. Hasan, S., Ahmed, M., Garcia-Ratés, S. & Greenfield, S. Antagonising a novel toxin “T14” in Alzheimer’s disease: Comparison of receptor blocker versus antibody effects in vitro. Biomed. Pharmacother. 158, 114120 (2023).

36. Szklarczyk, D. et al. STRING v11: protein-protein association networks with increased coverage, supporting functional discovery in genome-wide experimental datasets. Nucleic Acids Res. 47, D607–D613 (2019).

37. Kabbani, N. Proteomics of membrane receptors and signaling. PROTEOMICS 8, 4146–4155 (2008).

38. Brohée, S. & van Helden, J. Evaluation of clustering algorithms for protein-protein interaction networks. BMC Bioinformatics 7, 488 (2006).

39. Paraoanu, L. E. & Layer, P. G. Acetylcholinesterase in cell adhesion, neurite growth and network formation. FEBS J. 275, 618–624 (2008).

40. Greenfield, S. Discovering and targeting the basic mechanism of neurodegeneration: the role of peptides from the C-terminus of acetylcholinesterase: non-hydrolytic effects of ache: the actions of peptides derived from the C-terminal and their relevance to neurodegeneration. Chem. Biol. Interact. 203, 543–546 (2013).

41. Jean, L., Brimijoin, S. & Vaux, D. J. In vivo localization of human acetylcholinesterase-derived species in a β-sheet conformation at the core of senile plaques in Alzheimer’s disease. J. Biol. Chem. 294, 6253–6272 (2019).

42. Henley, J. & Poo, M. Guiding neuronal growth cones using Ca2+ signals. Trends Cell Biol. 14, 320–330 (2004).

43. Mingorance-Le Meur, A., Mohebiany, A. N. & O’Connor, T. P. Varicones and Growth Cones: Two Neurite Terminals in PC12 Cells. PLoS ONE 4, e4334 (2009).

44. Greenfield, S. A., Day, T., Mann, E. O. & Bermudez, I. A novel peptide modulates alpha7 nicotinic receptor responses: implications for a possible trophic-toxic mechanism within the brain. J. Neurochem. 90, 325–331 (2004).

45. Nordman, J. C. et al. Axon targeting of the alpha 7 nicotinic receptor in developing hippocampal neurons by Gprin1 regulates growth. J. Neurochem. 129, 649–662 (2014).

46. Zou, Z., Tao, T., Li, H. & Zhu, X. mTOR signaling pathway and mTOR inhibitors in cancer: progress and challenges. Cell Biosci. 10, 31 (2020).

47. Brunkard, J. O. Exaptive Evolution of Target of Rapamycin Signaling in Multicellular Eukaryotes. Dev. Cell 54, 142–155 (2020).

48. Hay, N. & Sonenberg, N. Upstream and downstream of mTOR. Genes Dev. 18, 1926–1945 (2004).

49. Witayateeraporn, W. et al. α7-Nicotinic acetylcholine receptor antagonist QND7 suppresses non-small cell lung cancer cell proliferation and migration via inhibition of Akt/mTOR signaling. Biochem. Biophys. Res. Commun. 521, 977–983 (2020).

50. Kosciuczuk, E. M., Saleiro, D. & Platanias, L. C. Dual targeting of eIF4E by blocking MNK and mTOR pathways in leukemia. Cytokine 89, 116–121 (2017).

51. Majeed, S. T. et al. mTORC1 induces eukaryotic translation initiation factor 4E interaction with TOS-S6 kinase 1 and its activation. Cell Cycle Georget. Tex 20, 839–854 (2021).

52. Deleyto-Seldas, N. & Efeyan, A. The mTOR–Autophagy Axis and the Control of Metabolism. Front. Cell Dev. Biol. 9, (2021).

53. Mizushima, N. & Yoshimori, T. How to interpret LC3 immunoblotting. Autophagy 3, 542–545 (2007).

54. Shen, D.-N., Zhang, L.-H., Wei, E.-Q. & Yang, Y. Autophagy in synaptic development, function, and pathology. Neurosci. Bull. 31, 416–426 (2015).

55. Morita, M. et al. mTOR coordinates protein synthesis, mitochondrial activity and proliferation. Cell Cycle Georget. Tex 14, 473–480 (2015).

56. Gergalova, G., Lykhmus, O., Komisarenko, S. & Skok, M. α7 nicotinic acetylcholine receptors control cytochrome c release from isolated mitochondria through kinase-mediated pathways. Int. J. Biochem. Cell Biol. 49, 26–31 (2014).

57. Knorr, D. Y., Georges, N. S., Pauls, S. & Heinrich, R. Acetylcholinesterase promotes apoptosis in insect neurons. Apoptosis Int. J. Program. Cell Death 25, 730–746 (2020).

58. Pereira, L. et al. A cellular and regulatory map of the cholinergic nervous system of C. elegans. eLife 4, e12432 (2015).

59. Phillis, J. W. Acetylcholine release from the central nervous system: a 50-year retrospective. Crit. Rev. Neurobiol. 17, 161–217 (2005).

60. Friedman, J. R. et al. Acetylcholine signaling system in progression of lung cancers. Pharmacol. Ther. 194, 222–254 (2019).

61. Zimmermann, M. Neuronal AChE splice variants and their non-hydrolytic functions: redefining a target of AChE inhibitors? Br. J. Pharmacol. 170, 953–967 (2013).

62. Luo, X., Lauwers, M., Layer, P. G. & Wen, C. Non-neuronal Role of Acetylcholinesterase in Bone Development and Degeneration. Front. Cell Dev. Biol. 8, 620543 (2020).

63. Xiang, Y.-Y., Dong, H., Yang, B. B., Macdonald, J. F. & Lu, W.-Y. Interaction of acetylcholinesterase with neurexin-1β regulates glutamatergic synaptic stability in hippocampal neurons. Mol. Brain 7, 15 (2014).

64. Duysen, E. G. & Lockridge, O. Phenotype comparison of three acetylcholinesterase knockout strains. J. Mol. Neurosci. MN 30, 91–92 (2006).

65. Layer, P. G. & Willbold, E. Novel Functions of Cholinesterases in Development, Physiology and Disease. Prog. Histochem. Cytochem. 29, III–92 (1994).

66. Holmes, C., Jones, S. A., Budd, T. C. & Greenfield, S. A. Non-cholinergic, trophic action of recombinant acetylcholinesterase on mid-brain dopaminergic neurons. J. Neurosci. Res. 49, 207–218 (1997).

67. Brai, E., Stuart, S., Badin, A.-S. & Greenfield, S. A. A Novel Ex Vivo Model to Investigate the Underlying Mechanisms in Alzheimer’s Disease. Front. Cell. Neurosci. 11, (2017).

68. Liu, E. Y. L. et al. Interacting with α7 nAChR is a new mechanism for AChE to enhance the inflammatory response in macrophages. Acta Pharm. Sin. B 10, 1926–1942 (2020).

69. Broide, R. S., Robertson, R. T. & Leslie, F. M. Regulation of α7 Nicotinic Acetylcholine Receptors in the Developing Rat Somatosensory Cortex by Thalamocortical Afferents. J. Neurosci. 16, 2956–2971 (1996).

70. Laplante, M. & Sabatini, D. M. mTOR signaling in growth control and disease. Cell 149, 274–293 (2012).

71. Wullschleger, S., Loewith, R. & Hall, M. N. TOR signaling in growth and metabolism. Cell 124, 471–484 (2006).

72. Lv, G. et al. Inhibiting specificity protein 1 attenuated sevoflurane-induced mitochondrial stress and promoted autophagy in hippocampal neurons through PI3K/Akt/mTOR and α7-nAChR signaling. Neurosci. Lett. 794, 136995 (2023).

73. Ito, T. et al. The neuroprotective effects of activated α7 nicotinic acetylcholine receptor against mutant copper-zinc superoxide dismutase 1-mediated toxicity. Sci. Rep. 10, 22157 (2020).

74. Eleftheriadis, T., Pissas, G., Liakopoulos, V. & Stefanidis, I. Cytochrome c as a Potentially Clinical Useful Marker of Mitochondrial and Cellular Damage. Front. Immunol. 7, (2016).

75. Mueed, Z. et al. Tau and mTOR: The Hotspots for Multifarious Diseases in Alzheimer’s Development. Front. Neurosci. 12, 1017 (2019).

76. Subramanian, A. et al. Trilateral association of autophagy, mTOR and Alzheimer’s disease: Potential pathway in the development for Alzheimer’s disease therapy. Front. Pharmacol. 13, (2022).

77. Torres, M. et al. Defective lysosomal proteolysis and axonal transport are early pathogenic events that worsen with age leading to increased APP metabolism and synaptic Abeta in transgenic APP/PS1 hippocampus. Mol. Neurodegener. 7, 59 (2012).

78. Caccamo, A., Majumder, S., Richardson, A., Strong, R. & Oddo, S. Molecular interplay between mammalian target of rapamycin (mTOR), amyloid-beta, and Tau: effects on cognitive impairments. J. Biol. Chem. 285, 13107–13120 (2010).

79. Auld, D. S., Kornecook, T. J., Bastianetto, S. & Quirion, R. Alzheimer’s disease and the basal forebrain cholinergic system: relations to beta-amyloid peptides, cognition, and treatment strategies. Prog. Neurobiol. 68, 209–245 (2002).

