## Supplemental Table 1 for "The human acetylcholinesterase c-terminal T30 peptide activates neural growth through an alpha 7 nicotinic acetylcholine receptor mTOR pathway"

**Supplemental Table 1:** A complete list of significantly altered proteins and within SH-SY5Y cell treated with 100nM T30 for 3 DIV. Peptide abundance determined by precursor ion quantification in Proteome Discoverer v2.4.

| Gene Symbol | Description | $log_2$ fold change T30 | fold change T30 p-value |
|---|---|---|---|
| A2M | alpha-2-macroglobulin isoform X1 | -2.41E+00 | 1.02E-07 |
| AKT3 | RAC-gamma serine/threonine-protein kinase isoform X1 | 7.74E-01 | 3.72E-02 |
| ALB | serum albumin preproprotein | -1.19E+00 | 4.18E-03 |
| ANKRD44 | serine/threonine-protein phosphatase 6 regulatory ankyrin repeat subunit B isoform X1 | 1.68E+00 | 3.46E-03 |
| ARF5 | ADP-ribosylation factor 5 | 8.42E-01 | 6.39E-05 |
| ARHGAP33 | rho GTPase-activating protein 33 isoform X9 | -1.99E+00 | 3.22E-03 |
| ARL8A | ADP-ribosylation factor-like protein 8A isoform 1 | 7.03E-01 | 6.96E-04 |
| ARMC6 | armadillo repeat-containing protein 6 isoform 1 | 8.75E-01 | 8.40E-03 |
| ATP2B1 | plasma membrane calcium-transporting ATPase 1 isoform X1 | 1.35E+00 | 3.76E-04 |
| ATP5IF1 | ATPase inhibitor, mitochondrial isoform 1 precursor | -2.23E+00 | 2.16E-04 |
| ATP6V1C2 | V-type proton ATPase subunit C 2 isoform X1 | -9.05E-01 | 8.44E-03 |
| ATP6V1G1 | V-type proton ATPase subunit G 1 | 7.43E-01 | 2.71E-02 |
| BAG2 | BAG family molecular chaperone regulator 2 | 1.50E+00 | 4.07E-07 |
| BCAP31 | B-cell receptor-associated protein 31 isoform a | -1.46E+00 | 3.15E-02 |
| BCL2L13 | bcl-2-like protein 13 isoform X1 | 1.20E+00 | 7.46E-05 |
| BTF3L4 | transcription factor BTF3 homolog 4 isoform 1 | 6.85E-01 | 1.06E-03 |
| CAMK1G | calcium/calmodulin-dependent protein kinase type 1G | 9.85E-01 | 2.33E-07 |
| CENPF | centromere protein F | 1.50E+00 | 3.46E-02 |
| CHGB | secretogranin-1 precursor | 7.98E-01 | 3.58E-02 |
| CISD3 | CDGSH iron-sulfur domain-containing protein 3, mitochondrial precursor | 1.24E+00 | 1.65E-05 |
| CKS2 | cyclin-dependent kinases regulatory subunit 2 | 2.28E+00 | 5.11E-10 |
| CLUH | clustered mitochondria protein homolog isoform X1 | 1.03E+00 | 1.99E-07 |
| COPG1 | coatomer subunit gamma-1 | 7.27E-01 | 3.05E-02 |
| COTL1 | coactosin-like protein | 1.44E+00 | 4.69E-02 |
| COX5A | cytochrome c oxidase subunit 5A, mitochondrial precursor | -9.86E-01 | 6.22E-03 |
| CRADD | death domain-containing protein CRADD isoform 1 | 8.00E+00 | 1.36E-02 |

| | | | |
|---|---|---|---|
| CUTA | protein CutA isoform 1 | 5.68E-01 | 8.34E-03 |
| DENND2A | DENN domain-containing protein 2A isoform a | 1.21E+00 | 2.56E-03 |
| DEPDC1 | DEP domain-containing protein 1A isoform a | 1.23E+00 | 8.15E-05 |
| DHRS7B | dehydrogenase/reductase SDR family member 7B isoform X1 | 1.69E+00 | 5.08E-03 |
| DHTKD1 | probable 2-oxoglutarate dehydrogenase E1 component DHKTD1, mitochondrial | 2.88E+00 | 3.07E-16 |
| DLL3 | delta-like protein 3 isoform 1 precursor | 1.07E+00 | 7.70E-07 |
| DNAAF5 | dynein assembly factor 5, axonemal | 7.08E-01 | 1.81E-02 |
| DNAH10 | dynein heavy chain 10, axonemal isoform X1 | 3.34E+00 | 3.07E-16 |
| DNAJA4 | dnaJ homolog subfamily A member 4 isoform 1 | 1.26E+00 | 8.46E-10 |
| DNAJB4 | dnaJ homolog subfamily B member 4 isoform a | 1.07E+00 | 1.35E-02 |
| DPF2 | zinc finger protein ubi-d4 isoform X1 | 1.40E+00 | 3.40E-04 |
| EIF4ENIF1 | eukaryotic translation initiation factor 4E transporter isoform X1 | 1.73E+00 | 2.26E-03 |
| ENOPH1 | enolase-phosphatase E1 isoform 1 | 6.83E-01 | 1.06E-03 |
| ENY2 | transcription and mRNA export factor ENY2 isoform 1 | 8.38E-01 | 1.08E-04 |
| EYS | protein eyes shut homolog isoform 4 precursor | 1.87E+00 | 9.77E-03 |
| FAM213A | redox-regulatory protein FAM213A isoform X1 | 6.20E-01 | 2.75E-03 |
| FAM3C | protein FAM3C isoform X1 | -1.30E+00 | 9.81E-03 |
| FARS2 | phenylalanine--tRNA ligase, mitochondrial | 1.49E+00 | 1.03E-03 |
| FKBP15 | FK506-binding protein 15 | 1.33E+00 | 5.59E-04 |
| FLJ10769 | ATP-dependent (S)-NAD(P)H-hydrate dehydratase isoform a | 1.32E+00 | 1.04E-03 |
| FLOT2 | flotillin-2 isoform X1 | 1.07E+00 | 8.91E-03 |
| FSD1L | FSD1-like protein isoform X1 | 2.15E+00 | 8.46E-10 |
| FSIP2 | fibrous sheath-interacting protein 2 | -1.09E+00 | 2.11E-02 |
| GGA2 | ADP-ribosylation factor-binding protein GGA2 | 1.82E+00 | 5.18E-03 |
| GNE | bifunctional UDP-N-acetylglucosamine 2-epimerase/N-acetylmannosamine kinase isoform 1 | 9.47E-01 | 2.44E-02 |
| GOLGA4 | golgin subfamily A member 4 isoform X1 | 1.76E+00 | 5.33E-03 |
| GTF2A1 | transcription initiation factor IIA subunit 1 isoform 1 | 1.19E+00 | 3.01E-03 |
| H2AC6 | histone H2A type 1-C | 6.43E-01 | 2.92E-03 |
| H3C1 | histone H3.1 | -1.01E+00 | 1.31E-03 |
| HAUS8 | HAUS augmin-like complex subunit 8 isoform a | 1.71E+00 | 4.00E-03 |
| HBS1L | HBS1-like protein isoform 1 | 1.46E+00 | 1.66E-03 |
| HECTD4 | probable E3 ubiquitin-protein ligase HECTD4 | 1.07E+00 | 7.07E-03 |

| | | | |
|---|---|---|---|
| HEXA | beta-hexosaminidase subunit alpha isoform 1 precursor | 1.54E+00 | 3.11E-09 |
| HMCN1 | hemicentin-1 precursor | 2.58E+00 | 3.07E-16 |
| HMGN2 | non-histone chromosomal protein HMG-17 | -1.63E+00 | 1.83E-02 |
| HNRNPR | heterogeneous nuclear ribonucleoprotein R isoform X1 | 2.53E+00 | 3.85E-11 |
| IQSEC1 | IQ motif and SEC7 domain-containing protein 1 isoform X2 | 1.39E+00 | 1.34E-03 |
| IRAK1 | interleukin-1 receptor-associated kinase 1 isoform 1 | 2.50E+00 | 2.76E-05 |
| IRF3 | interferon regulatory factor 3 isoform X2 | 1.89E+00 | 3.51E-04 |
| ISOC2 | isochorismatase domain-containing protein 2 isoform 2 | 9.44E-01 | 7.24E-03 |
| ISY1 | ISY1-RAB43 protein | 9.72E-01 | 4.11E-04 |
| KBTBD3 | kelch repeat and BTB domain-containing protein 3 isoform 1 | -1.09E+00 | 4.35E-02 |
| KCNAB2 | voltage-gated potassium channel subunit beta-2 isoform X5 | 1.96E+00 | 1.89E-03 |
| KHDRBS1 | KH domain-containing, RNA-binding, signal transduction-associated protein 1 isoform 1 | -1.18E+00 | 2.48E-04 |
| KIF15 | kinesin-like protein KIF15 isoform X1 | 1.12E+00 | 1.72E-02 |
| KIF2C | kinesin-like protein KIF2C isoform 1 | 3.30E+00 | 3.07E-16 |
| LAMB1 | laminin subunit beta-1 isoform X1 | 1.66E+00 | 1.47E-05 |
| LAMTOR3 | ragulator complex protein LAMTOR3 isoform 1 | 5.50E-01 | 4.50E-02 |
| LMAN2 | vesicular integral-membrane protein VIP36 precursor | -8.97E-01 | 1.17E-02 |
| LSM8 | LSM8 homolog, U6 small nuclear RNA associated | -1.10E+00 | 4.70E-03 |
| LTN1 | E3 ubiquitin-protein ligase listerin isoform 1 | 1.23E+00 | 2.55E-03 |
| MAPK1 | mitogen-activated protein kinase 1 | -1.07E+00 | 1.88E-02 |
| MCUR1 | mitochondrial calcium uniporter regulator 1 | -8.60E-01 | 3.46E-02 |
| METTL16 | U6 small nuclear RNA (adenine-(43)-N(6))-methyltransferase | 1.64E+00 | 1.18E-02 |
| METTL3 | N6-adenosine-methyltransferase catalytic subunit | 5.18E-01 | 7.73E-03 |
| MRPL50 | 39S ribosomal protein L50, mitochondrial | 6.33E-01 | 3.22E-03 |
| MRPS35 | 28S ribosomal protein S35, mitochondrial isoform 1 precursor | 9.77E-01 | 7.20E-05 |
| MSI2 | RNA-binding protein Musashi homolog 2 isoform X1 | 1.10E+00 | 9.06E-08 |
| NAGA | alpha-N-acetylgalactosaminidase isoform X1 | 2.22E+00 | 2.40E-04 |
| NCOR1 | nuclear receptor corepressor 1 isoform X1 | 7.71E-01 | 2.01E-02 |
| NDE1 | nuclear distribution protein nudE homolog 1 isoform X5 | 1.42E+00 | 3.84E-04 |

| | | | |
|---|---|---|---|
| NDUFAF4 | NADH dehydrogenase [ubiquinone] 1 alpha subcomplex assembly factor 4 | 2.32E+00 | 6.08E-11 |
| NEDD8 | NEDD8 precursor | 7.11E-01 | 2.14E-04 |
| NF1 | neurofibromin isoform 1 | -1.27E+00 | 1.50E-02 |
| NME1 | nucleoside diphosphate kinase A isoform a | -1.09E+00 | 4.17E-02 |
| NMRAL1 | nmrA-like family domain-containing protein 1 isoform X2 | 1.64E+00 | 3.07E-16 |
| NMU | neuromedin-U isoform 1 preproprotein | 1.67E+00 | 7.95E-03 |
| NOC3L | nucleolar complex protein 3 homolog | 7.92E-01 | 1.38E-03 |
| NOL8 | nucleolar protein 8 isoform X1 | 9.15E-01 | 4.83E-02 |
| NOSIP | nitric oxide synthase-interacting protein isoform X3 | 7.65E-01 | 2.36E-02 |
| NT5DC1 | 5'-nucleotidase domain-containing protein 1 | 1.03E+00 | 2.38E-02 |
| NUCB1 | nucleobindin-1 isoform X1 | 4.09E-01 | 3.94E-02 |
| NVL | nuclear valosin-containing protein-like isoform X2 | 2.41E+00 | 1.56E-04 |
| NXN | nucleoredoxin isoform 1 | 1.37E+00 | 1.41E-02 |
| OPA1 | dynamin-like 120 kDa protein, mitochondrial isoform 8 | 5.50E-01 | 1.62E-02 |
| ORC2 | origin recognition complex subunit 2 isoform X1 | 2.00E+00 | 4.11E-04 |
| PLRG1 | pleiotropic regulator 1 isoform 1 | -9.49E-01 | 2.84E-02 |
| PODXL2 | podocalyxin-like protein 2 precursor | 1.98E+00 | 1.31E-03 |
| PPFIA1 | liprin-alpha-1 isoform X8 | 9.99E-01 | 6.38E-04 |
| PTMA | prothymosin alpha isoform X1 | -1.46E+00 | 4.98E-02 |
| PTPN2 | tyrosine-protein phosphatase non-receptor type 2 isoform X1 | 1.87E+00 | 2.75E-10 |
| PUS1 | tRNA pseudouridine synthase A isoform 1 | 1.52E+00 | 9.33E-05 |
| RAB11FIP1 | rab11 family-interacting protein 1 isoform 3 | 1.21E+00 | 9.66E-03 |
| RFC2 | replication factor C subunit 2 isoform 1 | 8.10E-01 | 1.96E-02 |
| RIPK1 | receptor-interacting serine/threonine-protein kinase 1 isoform 1 | 1.50E+00 | 3.36E-02 |
| RPL15 | 60S ribosomal protein L15 isoform 1 | -8.62E-01 | 1.99E-02 |
| RPL26L1 | 60S ribosomal protein L26-like 1 | -1.70E+00 | 3.07E-02 |
| RPLP1 | 60S acidic ribosomal protein P1 isoform 1 | 5.70E-01 | 3.13E-03 |
| RPS27 | 40S ribosomal protein S27 isoform 1 | -1.09E+00 | 4.92E-03 |
| RPS27A | ubiquitin-40S ribosomal protein S27a precursor | 4.93E-01 | 1.19E-02 |
| RPS5 | 40S ribosomal protein S5 | -8.78E-01 | 1.54E-02 |
| RSBN1L | round spermatid basic protein 1-like protein | 1.07E+00 | 1.62E-02 |
| RSU1 | ras suppressor protein 1 isoform 1 | 1.62E+00 | 1.44E-05 |
| S100A6 | protein S100-A6 | 1.13E+00 | 4.09E-03 |

| | | | |
|---|---|---|---|
| SCAF4 | splicing factor, arginine/serine-rich 15 isoform 1 | -1.29E+00 | 2.23E-03 |
| SCOC | short coiled-coil protein isoform 1 | 1.40E+00 | 3.51E-04 |
| SEC22B | vesicle-trafficking protein SEC22b precursor | 5.18E-01 | 2.70E-02 |
| SETD3 | histone-lysine N-methyltransferase setd3 isoform X1 | 1.17E+00 | 3.36E-03 |
| SGF29 | SAGA-associated factor 29 | 2.03E+00 | 1.12E-08 |
| SLC12A9 | solute carrier family 12 member 9 isoform X1 | 1.80E+00 | 1.31E-03 |
| SPG7 | paraplegin isoform X1 | 2.67E+00 | 3.07E-16 |
| SPIRE1 | protein spire homolog 1 isoform X1 | 1.73E+00 | 2.36E-02 |
| SRSF11 | serine/arginine-rich splicing factor 11 isoform 3 | 4.83E-01 | 3.21E-02 |
| STARD7 | stAR-related lipid transfer protein 7, mitochondrial precursor | 1.30E+00 | 2.53E-05 |
| STK38 | serine/threonine-protein kinase 38 isoform X1 | 5.33E-01 | 1.45E-02 |
| STUB1 | E3 ubiquitin-protein ligase CHIP isoform a | 5.15E-01 | 1.45E-02 |
| SUMO2 | small ubiquitin-related modifier 2 isoform a precursor | -1.16E+00 | 1.23E-04 |
| TAF1 | transcription initiation factor TFIID subunit 1 isoform X1 | 1.15E+00 | 7.02E-04 |
| TECPR1 | tectonin beta-propeller repeat-containing protein 1 | 1.39E+00 | 4.86E-02 |
| THOC2 | THO complex subunit 2 isoform X1 | 1.20E+00 | 2.00E-03 |
| TLR7 | toll-like receptor 7 precursor | -1.91E+00 | 2.97E-03 |
| TMA7 | translation machinery-associated protein 7 isoform 1 | -1.58E+00 | 7.11E-05 |
| TMEM167A | protein kish-A precursor | 1.42E+00 | 4.56E-02 |
| TMEM230 | transmembrane protein 230 isoform X1 | -1.47E+00 | 4.36E-02 |
| TMSB10 | thymosin beta-10 | -1.31E+00 | 1.04E-03 |
| TOMM40 | mitochondrial import receptor subunit TOM40 homolog | 1.32E+00 | 4.79E-02 |
| TP53BP1 | TP53-binding protein 1 isoform 1 | 2.77E+00 | 9.04E-07 |
| TP53RK | TP53-regulating kinase | 1.48E+00 | 3.85E-02 |
| TPM3 | tropomyosin alpha-3 chain isoform Tpm3.2cy | 2.26E+00 | 3.07E-16 |
| TXN2 | thioredoxin, mitochondrial isoform X1 | 1.92E+00 | 5.16E-04 |
| UBE2A | ubiquitin-conjugating enzyme E2 A isoform 1 | -1.50E+00 | 4.23E-02 |
| UBE3C | ubiquitin-protein ligase E3C | 6.67E-01 | 2.65E-02 |
| UQCRFS1 | cytochrome b-c1 complex subunit Rieske, mitochondrial | 1.24E+00 | 6.33E-05 |
| USP9X | probable ubiquitin carboxyl-terminal hydrolase FAF-X isoform X1 | 5.55E-01 | 2.50E-02 |
| UTP14A | U3 small nucleolar RNA-associated protein 14 homolog A isoform 1 | 1.68E+00 | 1.41E-02 |

| | | | |
|---|---|---|---|
| WDR3 | WD repeat-containing protein 3 | 2.27E+00 | 5.77E-04 |
| XPNPEP3 | probable Xaa-Pro aminopeptidase 3 isoform 1 | -8.65E-01 | 1.85E-02 |
| YBX1 | nuclease-sensitive element-binding protein 1 | -8.55E-01 | 3.63E-02 |
| ZDHHC6 | palmitoyltransferase ZDHHC6 isoform X6 | -2.44E+00 | 1.39E-04 |
| ZNF48 | zinc finger protein 48 isoform 1 | 1.29E+00 | 3.31E-03 |
| ZNF830 | zinc finger protein 830 | 1.63E+00 | 1.61E-02 |
